# A SARS-CoV-2 spike ferritin nanoparticle vaccine protects against heterologous challenge with B.1.1.7 and B.1.351 virus variants in Syrian golden hamsters

**DOI:** 10.1101/2021.06.16.448525

**Authors:** Kathryn McGuckin Wuertz, Erica K. Barkei, Wei-Hung Chen, Elizabeth J. Martinez, Ines Lakhal-Naouar, Linda L. Jagodzinski, Dominic Paquin-Proulx, Gregory D. Gromowski, Isabella Swafford, Akshaya Ganesh, Ming Dong, Xiankun Zeng, Paul V. Thomas, Rajeshwer S. Sankhala, Agnes Hajduczki, Caroline E. Peterson, Caitlin Kuklis, Sandrine Soman, Lindsay Wieczorek, Michelle Zemil, Alexander Anderson, Janice Darden, Heather Hernandez, Hannah Grove, Vincent Dussupt, Holly Hack, Rafael de la Barrera, Stasya Zarling, James F. Wood, Jeffrey W. Froude, Matthew Gagne, Amy R. Henry, Elham Bayat Mokhtari, Prakriti Mudvari, Shelly J. Krebs, Andrew S. Pekosz, Jeffrey R. Currier, Swagata Kar, Maciel Porto, Adrienne Winn, Kamil Radzyminski, Mark G. Lewis, Sandhya Vasan, Mehul Suthar, Victoria R. Polonis, Gary R. Matyas, Eli A. Boritz, Daniel C. Douek, Robert A. Seder, Sharon P. Daye, Mangala Rao, Sheila A. Peel, M. Gordon Joyce, Diane L. Bolton, Nelson L. Michael, Kayvon Modjarrad

## Abstract

The emergence of SARS-CoV-2 variants of concern (VOC) requires adequate coverage of vaccine protection. We evaluated whether a spike ferritin nanoparticle vaccine (SpFN), adjuvanted with the Army Liposomal Formulation QS21 (ALFQ), conferred protection against the B.1.1.7 and B.1.351 VOCs in Syrian golden hamsters. SpFN-ALFQ was administered as either single or double-vaccination (0 and 4 week) regimens, using a high (10 μg) or low (0.2 μg) immunogen dose. Animals were intranasally challenged at week 11. Binding antibody responses were comparable between high- and low-dose groups. Neutralizing antibody titers were equivalent against WA1, B.1.1.7, and B.1.351 variants following two high dose two vaccinations. SpFN-ALFQ vaccination protected against SARS-CoV-2-induced disease and viral replication following intranasal B.1.1.7 or B.1.351 challenge, as evidenced by reduced weight loss, lung pathology, and lung and nasal turbinate viral burden. These data support the development of SpFN-ALFQ as a broadly protective, next-generation SARS-CoV-2 vaccine.

## Introduction

Since the beginning of the COVID-19 pandemic, an unprecedented global effort has resulted in the rapid emergence, and early regulatory authorization or approval, of multiple vaccines for use in humans^1^. The emergence of variants of concern (VOC) underscores the need for continued development of next-generation vaccines^2,3^. Increasingly, VOCs are becoming the dominant circulating lineages world-wide, owing to their increased transmissibility and potential to cause breakthrough infection in vaccinated individuals. Two VOCs have gained particular attention: B.1.1.7 and B.1.351, first detected in the United Kingdom and in the Republic of South Africa, respectively^4–10^. Recent studies have shown markedly reduced cross-neutralizing antibody responses from convalescent and vaccinee sera against both VOCs, but B.1.351 in particular ^4,5,8,11–14^. Consequently, there has been a renewed, global effort to new iterations of SARS-CoV-2 vaccines confer protection against current and emerging SARS-CoV-2 VOCs^4,15,16^.

We recently developed a novel SARS-CoV-2 vaccine that presents eight prefusion-stabilized spike glycoprotein trimers in an ordered array on a ferritin nanoparticle (SpFN) and adjuvanted with Army Liposomal Formulation QS21 (ALFQ). This formulation has demonstrated efficacy against the origin strain of SARS-CoV-2 (WA1) in both K18 murine and rhesus macaque viral challenge models^17–20^. In rhesus macaques, SpFN elicited a dose-dependent potent humoral and cellular immune response that translated into a precipitous reduction in viral load upper and lower airways of the animals, as well as protection from lung histopathology. Importantly, immunization induced cross-neutralizing antibodies against current circulating VOCs^18^. Building on these data, we evaluated the efficacy of SpFN-ALFQ to protect against virus challenge with VOCs B.1.1.7 and B.1.351 in a Syrian golden hamster (SGH) challenge model. This animal model has become a standard in the field for pre-clinical SARS-CoV-2 vaccine development, as respiratory pathology in this model closely recapitulates human disease^21–24^.

Here, we demonstrate that SpFN adjuvanted with ALFQ (SpFN-ALFQ) generates strong binding antibody responses against the receptor binding domain and spike proteins of both B.1.1.7 and B.1.351, as well as potent neutralizing antibody responses against both VOCs. Consistent with development of a protective humoral response, we demonstrate that SpFN-ALFQ confers clear protection against upper respiratory tract disease following challenge with these VOCs, as demonstrated by less body weight loss and decreased tissue viral burden and lung pathology. The SpFN vaccine candidate is now currently under assessment in a human phase 1 clinical trial (ClinicalTrials.gov Identifier: NCT04784767). These data support further development of the vaccine as one that may be broadly applicable to multiple sarbecovirus lineages.

## Results

### Hamster model to assess efficacy of SpFN-ALFQ against B.1.1.7 and B.1.351

Prior studies have demonstrated that the SpFN-ALFQ vaccine is efficacious against the WA1 strain of SARS-CoV-2 in a nonhuman primate model^18^. The ability of this vaccine to confer protection against the VOC B.1.1.7, now prominent throughout the U.S. and other parts of the world, and B.1.351, prominent in Africa and emerging world-wide, is unknown. We evaluated vaccine immunogenicity and efficacy against VOCs in SGH due to the susceptibility of SGH to severe clinical disease, lung pathology, and viral replication in the respiratory tract^24,25^. SpFN (Fig. 1a) adjuvanted with ALFQ was administered at a high (10 μg) or low dose (0.2 μg), selected based on immunogenicity and efficacy studies of SpFN-ALFQ in mice^20^, in a either a single (1) and two (2)-vaccination regimen in parallel with 2 injections of phosphate buffered saline (PBS) in control animals (Fig. 1b). Blood for serologic analysis was drawn periodically throughout the study, with key immunogenicity timepoints at weeks 6 and 8 (two and four weeks following last vaccine dose, respectively) and week 11 (seven weeks post final vaccine dose), preceding viral challenge.

**Figure 1.**
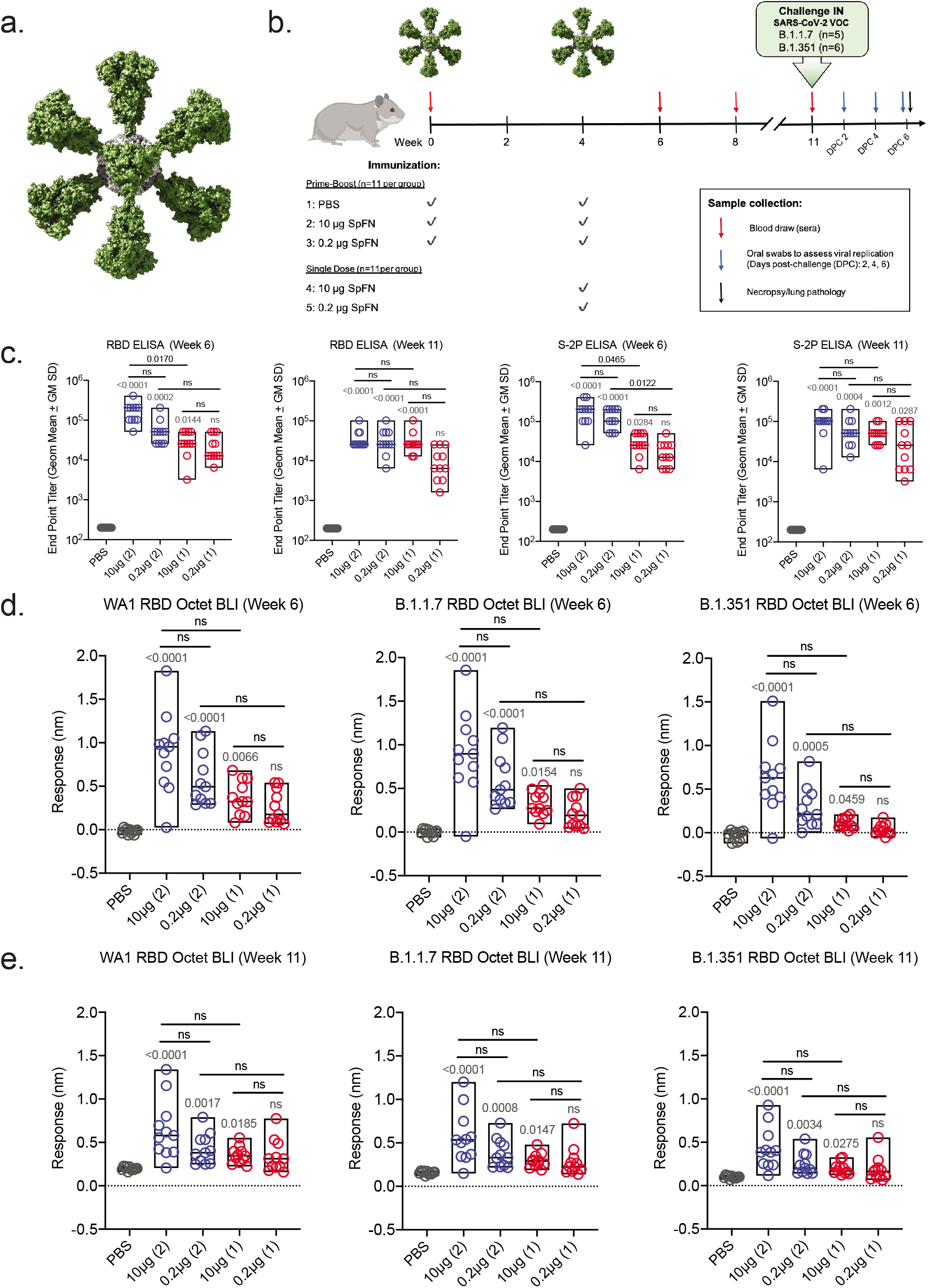
Antibody responses following SpFN-ALFQ immunization. **a,** The 3-dimensional model of SpFN with Spike protein trimers (green) decorating a ferritin core (gray) as viewed down a 3-fold axis. **b**, Experimental design with hamsters receiving immunization at weeks 0 and 4 in the 2-dose regimen and week 4 in the 1-dose regimen as depicted by green SpFN structures above the time-line and check marks below the timeline indicating immunogen dose (10 μg or 0.2 μg or PBS control). Phlebotomy samples were taken at weeks 0, 6, 8 and 11 as indicated by red arrows. Intranasal (IN) viral challenge was performed at week 11 with either VOC B.1.1.7 or B.1.351 viral stocks with the number of animals challenged noted parenthetically. Oral swabs were collected at 2, 4, and 6 days post challenge (DPC) as indicated by blue arrows above the timeline. Necropsy was performed on all animals at day 6 post-challenge as indicated by the black arrow. **c**, ELISA was performed using either WA1 derived Receptor Binding Domain (RBD) or S-2P Spike proteins from sera taken at weeks 6 and 11. Sera from week 0 was also assessed, with no detectable signal observed in any samples (data not shown). The vaccine regimens are indicated on the x-axis by PBS (control) or SpFN dose with the number of vaccinations in the regimen given parenthetically and by color code (blue, 2-dose, red, 1-dose). Endpoint titers are given on the y-axis as geometric mean titers with data displayed in box plots with the top and bottom bars of the box the standard deviation and the middle bar as the median value. P-values for active vaccination groups compared with PBS control are given just above the boxes in light grey while intra-active regimen p-values are given above the boxes in black. ns, not significant (p > 0.05) using the Kruskal-Wallis multiple comparisons test, with Dunn’s correction. **d,e**, Octet Biolayer Interferometry (BLI) responses against the WA1, B.1.1.7, and B.1.351 sequences of the RBD are given for the vaccination regimens as in C on the x-axis at weeks 6 (**d**) and 11 (**e**). BLI responses are given in nanometers (nm) on the y-axis. Color coding and statistical treatments are as in **c**.

### Strong binding antibody responses elicited by two-dose SpFN-ALFQ vaccination

Robust serum binding antibody responses to the SARS-CoV-2 WA1 spike protein (S-2P) and receptor-binding domain (RBD) were observed at week 6 by ELISA (Fig. 1c). Both the 1- and 2-dose regimens were immunogenic, with higher ELISA titers elicited by the 2-dose regimen against both RBD and spike (S-2P), while responses within regimens did not differ between doses. Endpoint titers against spike (S-2P) at week 6 demonstrated nearly a log difference between dosing regimens, with mean reciprocal dilutions for the 1-dose regimens of 3.2 × 10^4^ (10 μg) and 1.92 × 10^4^ (0.2 μg), and 2-dose regimen responses of 1.98 × 10^5^ (10 μg) and 1.35 × 10^5^ (0.2 μg). Endpoint titers did not substantially wane between weeks 6 (Fig. 1c) and 8 (Supplemental Fig. 1a) or week 11 (Fig. 1c), indicating durability of binding antibody responses.

We then assessed the breadth of post-vaccination binding antibody responses following to WA1 and VOCs B.1.1.7 and B.1.351 variant RBD antigens, measured by biolayer interferometry (Fig. 1d). Responses again trended higher with the 2-dose regimen and did not differ between the 10 μg and 0.2 μg groups within regimens. Binding levels between strains were comparable between WA1 and B.1.1.7 in the 2-dose regimens with B.1.1.7/WA1 mean fold change ratios of 1.02 (10 μg) and 1.01 (0.2 μg), while 1-dose group of B.1.1.7/WA1 ratios were 0.88 (10 μg) and 0.82 (0.2 μg). Comparatively, B.1.351 cross-reactive binding antibody levels were reduced compared to WA1. B.1.351/WA1 mean fold-change ratios in the 2-dose groups were 0.71 (10 μg) and 0.49 (0.2 μg), and in the 1-dose groups 0.29 (10 μg) and 0.14 (0.2 μg). Similar magnitude responses were observed at week 8 (Supplemental Fig. 1b), while binding antibody levels declined by week 11, the time of challenge (Fig 1e). Cross-reactive binding antibodies at week 6 also recognized heterologous B.1.1.7 and B.1.351 cell surface expressed spike protein in a flow cytometric based IgG opsonization assay; responses were consistent with the quantitative differences seen for the inter-group comparisons in Fig. 1 for all three strains (Supplemental Fig. 1c)

### hACE2 competition and pseudovirus neutralizing antibody responses

As SARS-CoV-2 mediates host cell entry via engagement of the viral spike protein with cell surface expressed human angiotensin-converting enzyme 2 (hACE2) protein, preventing this interaction is a critical target for vaccine induced immune responses^26^. We measured hACE2 inhibitory antibodies longitudinally using a serum competition assay that detects blockade of RBD-hACE2 binding (Fig. 2a). At week 6, mean 50% inhibitory dilution (ID_50_) values in the 2-dose regimen groups were 145.6 (10 μg) and 106.8 (0.2 μg), and 76.6 (10 μg) and 57.0 (0.2 μg) in the 1-dose groups. Responses did not differ between regimens and were largely stable at week 8 but waned by the time of challenge at week 11.

**Figure 2.**
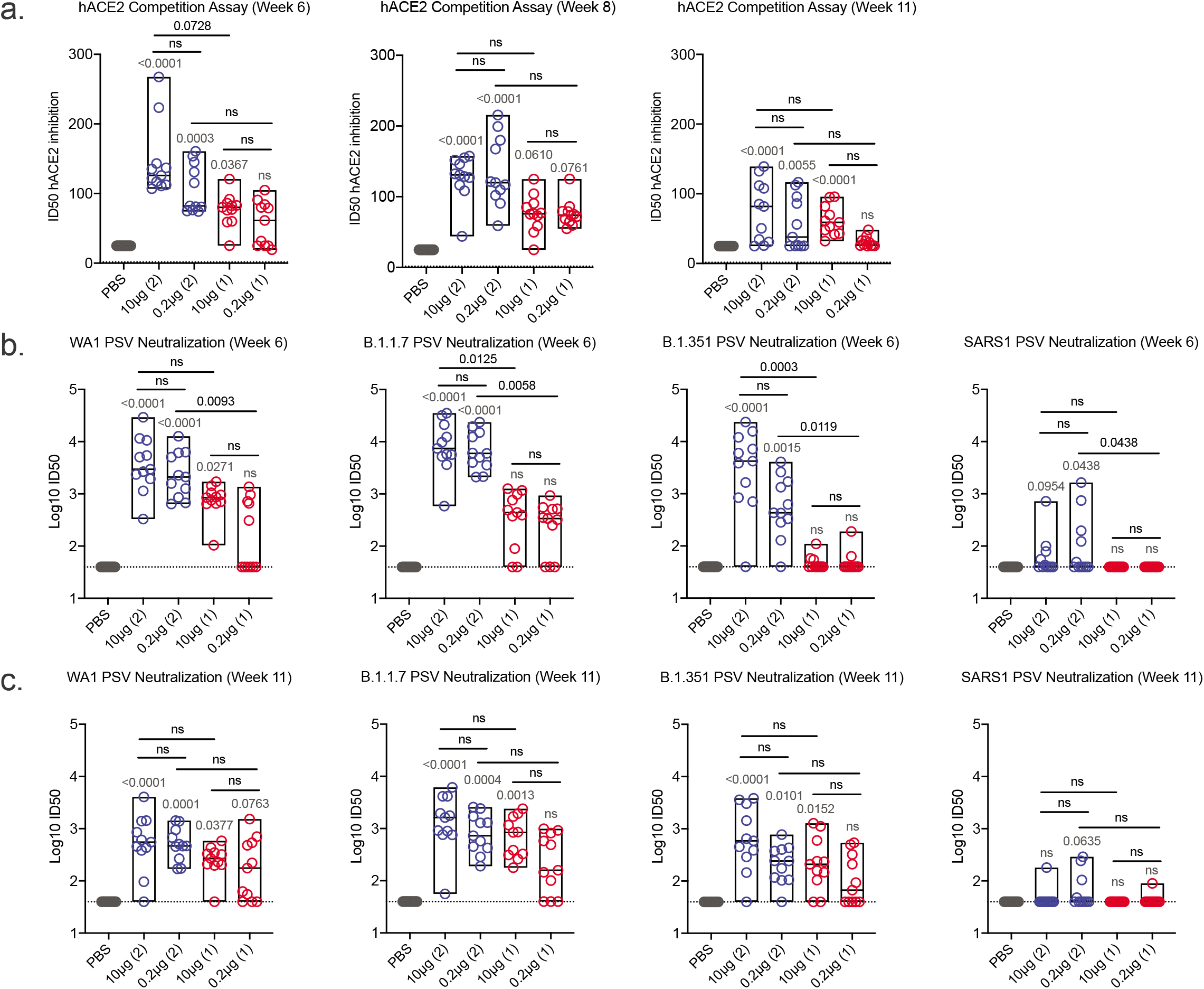
Human angiotensin-converting enzyme competition and pseudovirus neutralization responses following SpFN-ALFQ immunization. **a,** Human angiotensin-converting enzyme competition (hACE2) assays were performed from sera taken at week 6, 8, and 11. The vaccine regimens are indicated on the x-axis by PBS (control) or SpFN dose with the number of vaccinations in the regimen given parenthetically and by color code (blue, 2-dose, red, 1-dose). Inhibitory dose 50% (ID_50_) are given on the y-axis as with data displayed in box plots with the top and bottom bars of the box the standard deviation and the middle bar as the median value. P-values for SpFN-ALFQ vaccination groups compared with PBS control are given just above the boxes in light grey while intra-active regimen p-values are given above the boxes in black. ns, not significant (p > 0.05) using the Kruskal-Wallis multiple comparisons test, with Dunn’s correction. **b,c,** Pseudovirus neutralization at weeks 6 and 11 are given against Spike proteins derived from WA1, B.1.1.7, B.1.351 SARS-CoV-2 variants and from SARS-CoV-1. The vaccine regimens are indicated on the x-axis as in **a**. The neutralization titers are given as inhibitory dose 50% (ID_50_) on the y-axis which is a logarithmic scale. The data are given as box plots and statistically treated as described in **a**.

Pseudovirus neutralizing antibody responses were measured against the WA1, B.1.1.7 and B.1.351 variants at weeks 6 and 11 (Fig. 2b and Fig. 2c, respectively). At week 6, responses were higher for the 2-dose versus the 1-dose vaccine regimen in most cases (Fig. 2b). Mean log10 ID_50_ values against WA1 ranged from 3.56 (10 μg) to 3.36 (0.2 μg) and 2.88 (10 μg) to 2.17 (0.2 μg) to WA1 in the 2-dose and one-dose regimens, respectively, with comparable titers against B.1.1.7 of 3.90 (10 μg), 3.82 (0.2 μg), 2.52 (10 μg) and 2.36 (0.2 μg), respectively. Responses to B.1.351 were diminished relative to WA1 and B.1.1.7, notably with the one-dose regimen, with mean log10 ID_50_ values of 3.46 (10 μg) to 2.74 (0.2 μg) and 1.67 (10 μg) to 1.68 (0.2 μg) in the 2-dose and 1-dose regimens, respectively. Titers were consistently higher for the 2-dose versus the 1-dose vaccine regimen. Neutralizing antibody responses decreased between weeks 6 and 11 and the differences between the 2-dose and 1-dose regimens were less marked (Fig. 2c). Interestingly, cross-neutralization was observed for some animals in the 2-dose regimen against the related sarbecovirus SARS-CoV-1, albeit at reduced levels compared to SARS-CoV-2. Here we demonstrate that SpFN-ALFQ vaccine induces potent neutralization in SARS-CoV-2 VOCs, as well as cross-neutralization against SARS-CoV-1, as previously described in rhesus macaques and mice^18–20^.

### Reduction of body-weight loss in SpFN-ALFQ vaccinated, VOCs challenged hamsters

Hamsters were divided into two cohorts comprised of male and female animals and challenged with either B.1.1.7 (n=5) or B.1.351 (n=6) variants of SARS-CoV-2 by intranasal inoculation at week 11, 7 weeks after the last immunization in either the 2-dose or 1-dose vaccine regimen groups. Challenge doses were selected based on prior titration of viral stocks in hamsters, targeting a 10-15% loss of body weight by 7 days post-challenge. Body weight was monitored daily following challenge. For both B.1.1.7 and B.1.351 infection, weight loss of PBS control animals fell as expected between 10-15% by 6 days post-challenge (DPC), with a mean weight loss of 11.6% for B.1.1.7 and 12.7% for B.1.351 respectively (Fig. 3a). For the 10 μg 2-dose vaccinated animals, B.1.1.7 challenged animals had a mean body weight loss of 2.3% and B.1.351 challenged animals with slightly higher at 4.0%, but still a dramatic reduction from the PBS vaccinated control hamsters. In the 2-dose vaccine regimen groups, both the 10 μg and 0.2 μg dose groups demonstrated comparable levels of protection, with the one-dose vaccinated animals demonstrating the least protection both for B.1.1.7 and B.1.351 challenges.

**Figure 3.**
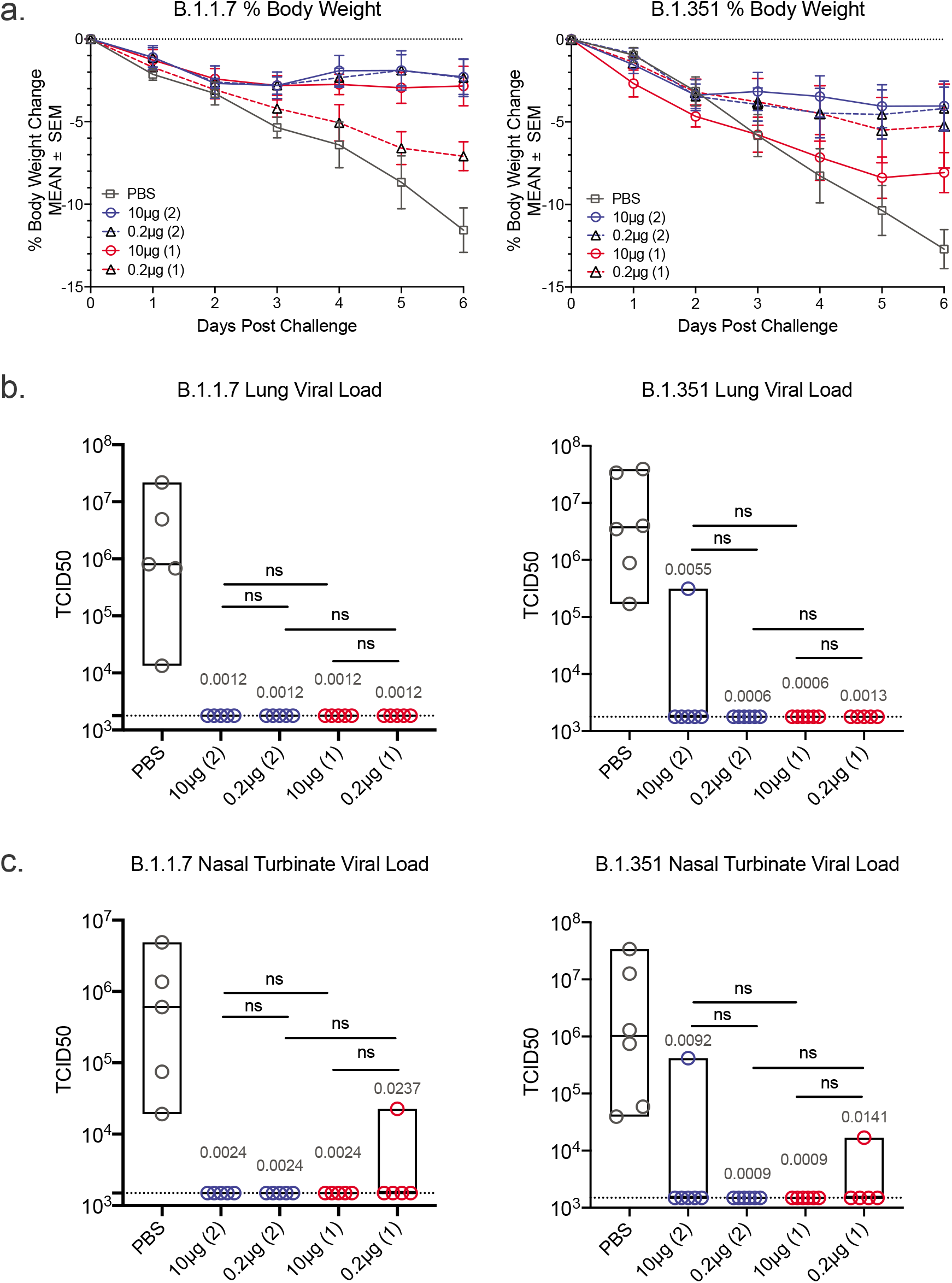
Body weight changes and lung viral load post-challenge. Daily weights were gathered on hamsters from the time of viral challenge to necropsy on day 6 post challenge when lungs were harvested for a whole tissue, culture based viral load assessment in Vero TMPRSS2 cells. **a**, Mean percent body weight changes plus and minus standard error of the mean (SEM) are given on the y-axis for groups of hamsters assigned to phosphate buffered saline control (PBS) or SpFN-ALFQ vaccination from day of challenge (day 0) to day of necropsy (day 6) for either B.1.1.7 or B.1.351 challenge. Data from immunization groups are given in each graph as PBS (grey plot, phosphate buffered saline control) or active immunogen dose (blue circle/ blue solid line, 10 μg 2-dose regimen; black triangle/ blue dotted line, 0.2 μg 2-dose regimen; red circle/red solid line, 10 μg 1-dose regimen; black triangle/red dotted line, 0.2 μg 1-dose regimen. Number of vaccinations in the vaccine regimen are also given parenthetically within each graph. **b**, SARS-CoV-2 viral load data from lung tissue harvested on day 6 post challenge is given on the y-axis as the tissue culture infective dose, 50% (TCID50) per gram of tissue as titered on Vero TMPRSS2 cells and read out by cytopathic effects for either B.1.1.7 and B.1.351 challenged hamsters. Immunization groups are given on the x-axis as PBS (phosphate buffered saline) control (gray circles); 10 μg and 0.2 μg 2-dose vaccine regimens (blue circles); 10 μg and 0.2 μg 1-dose vaccine regimens (red circles). Group data are plotted with the median group value given by the middle bar of the box plot. The dotted horizontal line is the lower limit of detection of the assay. **c,** SARS-CoV-2 viral load data from nasal turbinate tissue harvested on day 6 post challenge is given as in **b**. **b, c,** P-values for SpFN-ALFQ vaccination groups compared with PBS control are given just above the boxes in light grey while intra-active regimen p-values are given above the boxes in black. ns, not significant (p > 0.05) using the Kruskal-Wallis multiple comparisons test, with Dunn’s correction.

### Undetectable lung and nasal tissue viral load in SpFN-ALFQ vaccinated, viral challenged hamsters

To assess the impact of SpFN immunization following challenge by either the B.1.1.7 or B.1.351 strains, viral load was assessed by viral culture recovery in lung tissues at day 6 post challenge (Fig. 3b). The mean viral load in the PBS vaccinated control animals challenged with B.1.1.7 was 5.67 × 10^6^ TCID50/gram of lung tissue. Virus was not recovered in lung tissue culture in both the 10 and 0.2 μg 2-dose and 1-dose vaccine groups (below the limit of detection of 1.78 × 10^3^) (Fig. 3b). The mean lung tissue viral load in the PBS control vaccinated animals challenged with B.1.351 was 1.36 × 10^7^ TCID50/g. All animals in the 2-dose and 1-dose vaccine groups, for both 10 μg and 0.2 μg doses, showed no detectable virus except for one animal in the 10 μg two-dose vaccinated group (Fig. 3b). Viral load was also measured in nasal turbinate tissue collected at day 6 post-challenge, with similar results to lung tissue analysis (Fig. 3c). The mean nasal turbinate tissue viral load in the PBS control vaccinated animals challenged with B.1.1.7 was 1.39 × 10^6^ TCID50/g and B.1.351 was 8.14 × 10^6^ TCID50/g of nasal turbinate, respectively. All vaccinated animals were below the limit of detection (1.492 × 10^3^ TCID50/g), except a single outlier animal in the 0.2 μg 1-dose vaccination regimen group for B.1.1.7 challenged group, and an outlier animal each in the 10 μg 2-dose and 0.2 μg 1-dose regimen vaccination groups for B.1.351 challenged animals (Fig. 3c). Throughout the challenge phase of the study, oral swabs were collected at days 2, 4 and 6 post-challenge and assessed for viral burden by RT-qPCR, by measuring both sub-genomic E messenger RNA (sgmRNA) (Supplemental Fig. 2a and 2c) and total viral RNA (viral load) (Supplemental Fig. 2b and 2d). Overall viral loads decreased modestly over the duration of the challenge in all groups for both B.1.1.7 and B.1.351, with the largest decrease observed at day 6 post-challenge in the 2-dose 10 μg dose vaccination regimen group. These results demonstrate clear protection from tissue viral load in vaccinated animals challenged with B.1.1.7 and B.1.351, a critical determinant in establishing effective vaccines against SARS-CoV-2 VOCs.

### Lung pathology and tissue nucleocapsid antigen are minimized by SpFN-ALFQ vaccination

Lung pathology was assessed day 6 post-challenge by both routine hematoxylin and eosin (H&E) staining as well as immunohistochemistry (IHC) for the presence of viral nucleocapsid (N) protein (Fig. 4). By semiquantitative scoring of lung histopathology, the highest degree of pathology was seen in the PBS vaccinated control animals challenged with either B.1.1.7 or B.1.351 (Fig. 4a). All PBS vaccinated control animals developed histopathologic evidence of multifocal to extensive, moderate to marked interstitial pneumonia (IP) (Fig. 4b and 4c, for B.1.1.7 and B.1.351 challenged animals, respectively). The pneumonia was characterized by type II pneumocyte hyperplasia, alveolar edema, alveolar inflammatory and necrotic debris, thickening of alveolar septae, bronchiolar epithelial hyperplasia, and increased numbers of pulmonary macrophages (including multinucleated giant cells). The extent of IP present in PBS vaccinated control animals was comparable in animals challenged with either B.1.1.7 or B.1.351. In the B.1.1.7 challenged animals, the least amount of pathology was seen in the animals vaccinated with either 2-dose regimen, although less pathology was evident in all of the SpFN vaccination groups compared with the PBS control group (Fig. 4a - left panel, 4b, 4c and Supplemental Table 1A). For the B.1.351 challenged animals, the most protection from lung pathology was observed in the 10 μg, 2-dose vaccine regimen group, although there was a decrease in the mean pathology score of all SpFN vaccinated groups (Fig 4a - right panel, 4b, 4c and Supplemental Table 1A). Overall, there was a trend toward increased pathology observed in the B.1.351 versus the B.1.1.7 animals, in particular with the 1-dose groups. In the 10 μg 2-dose group however, protection between VOCs were comparable. Immunohistochemistry demonstrated strong, multifocal to focally extensive (>500-1000 cells per section) immunopositivity to SARS-CoV-2 nucleocapsid (N) protein in bronchiolar epithelium, alveolar pneumocytes and pulmonary macrophages in the lungs of all unvaccinated animals. Viral antigen detected in lung sections from animals in the 10 μg and 0.2 μg vaccine groups was substantially reduced compared with the PBS vaccinated groups with the greatest reduction seem in the 10 μg 2-shot vaccine groups (Supplemental Table 1B). Taken together, SpFN adjuvanted with ALFQ confers clear protection from viral burden and pathology in the lungs following VOC challenge.

**Figure 4.**
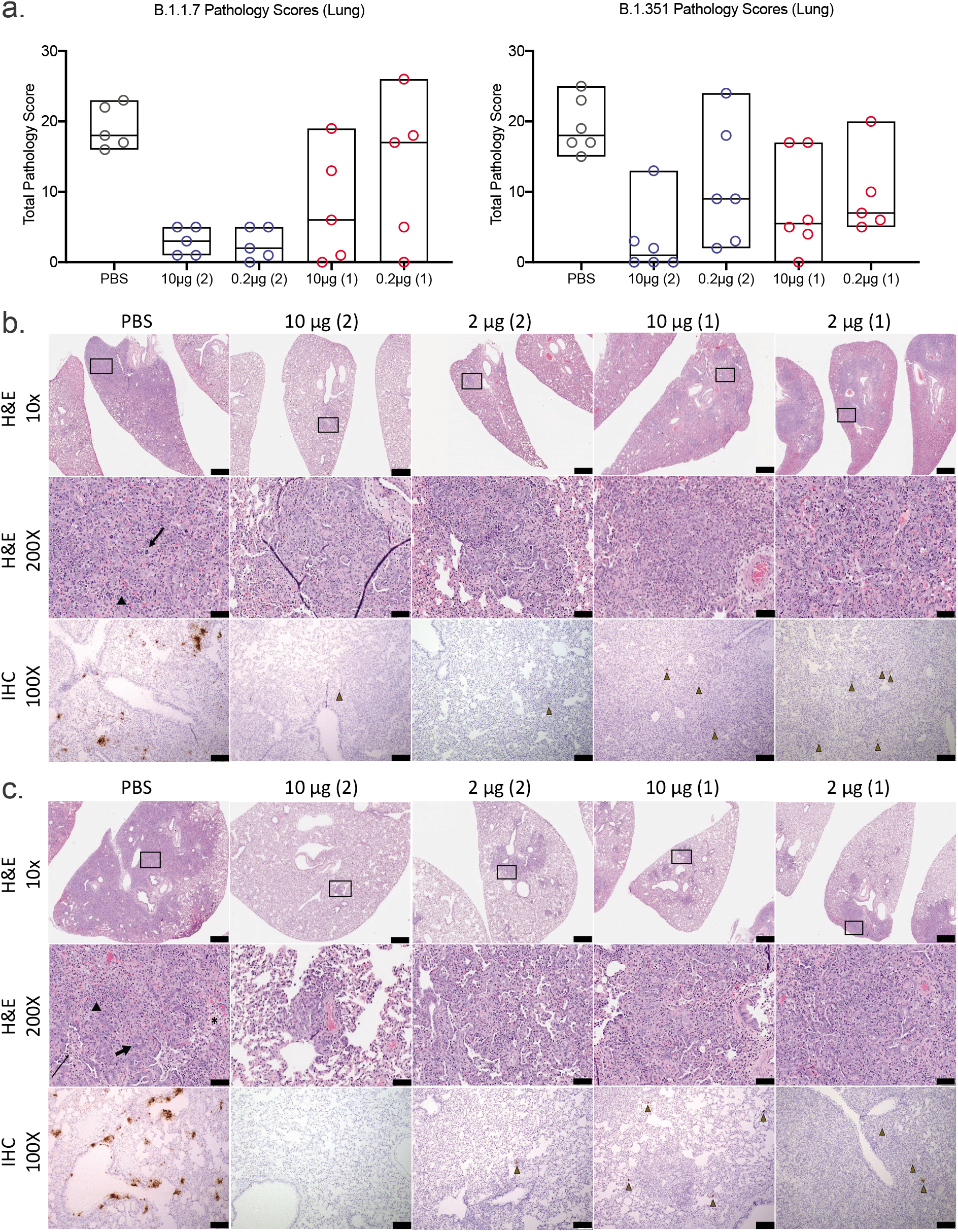
Standard and immunohistopathologic examination post-challenge. Lung tissues were collected at necropsy on day 6 post-challenge, fixed with neutral buffered formalin, and stained with hematoxylin and eosin (H&E) for standard microscopic examination as well has submitted for immunohistochemical (IHC) staining for SARS-CoV-2 nucleocapsid (N) protein. **a**, H&E stained slides were scored for pathologic effects on the y-axis (see Methods) for B.1.1.7 (left) and B.1.351 (right) challenged hamsters. Vaccination groups were plotted in box plots where the horizontal bar is the median group score. Vaccination groups are given as: PBS (phosphate buffered saline) control (gray circles); 10 μg and 0.2 μg 2-dose vaccine regimens (blue circles); 10 μg and 0.2 μg 1-dose vaccine regimens (red circles). **b,c,** Representative lung tissue sections from the PBS control and 10 μg and 0.2 μg 2-dose and 1-dose regimens with the number of vaccinations given parenthetically for B.1.1.7 (**b**) and B.1.351 (**c**) challenged hamsters in the columns as indicated. Rows are given by either H&E at 10- and 200-times magnification power (10X and 200X, respectively) or IHC of SARS-CoV-2 viral antigen at 100 times magnification power (100X). The black boxes in the top row indicate the area magnified in the middle row. Interstitial pneumonia is characterized by inflammatory cellular infiltrates (triangle), type II pneumocyte hyperplasia (thick arrow), bronchiolar epithelial hyperplasia, bronchiolar exudate (thin arrow) and edema (asterisk). SARS-CoV-2 immunopositive cells are highlighted by brown triangles. Scale bars: Top row, 1 mm; middle row, 50 μm; bottom row, 100 μm.

## Discussion

The development of effective and safe vaccines in response to the SARS-CoV-2 pandemic has occurred at an unprecedented rate. This success has been tempered by the reduced vaccine efficacy against some emerging VOCs, most notably B.1.351^4,5,8,11,13,16,27^. The need for more broadly effective vaccines against multiple lineages of SARS-CoV-2 will likely increase, as underscored by the rapid rise of VOCs in India, including B.1.1.7 and emerging variants B.1.617 with derivative lineages, and B.1.618^3^. As B.1.1.7 has become the dominant variant in the U.S.^28^ and B.1.351 and B.1.617 lineages are becoming more dominant in many areas of the world, evaluation of existing and novel vaccines against these strains is essential. As the pandemic continues, SARS-CoV-2 will not only adapt to replicate more efficiently in the human host and evolve to escape host immune responses to natural infection, it will increasingly adapt to vaccine evoked immune responses. This large number of products administered globally, with differential vaccine efficacy, will further drive the evolution of virus variants that may be more resistant to current formulations of SARS-CoV-2 vaccines. This will be especially true in populations where immunocompromised individuals are prevalent (e.g., HIV infection, cancer therapies, organ transplant, autoimmune diseases) providing an ideal milieu for the generation of virus variants to challenge current and next generation SARS-CoV-2 vaccines^29^.

We evaluated the efficacy of a novel vaccine, currently in clinical trials (ClinicalTrials.gov Identifier: NCT04784767), against VOCs. SpFN adjuvanted with ALFQ, has been previously demonstrated to be highly efficacious against the WA1 strain of SARS-CoV-2 in nonhuman primate and murine models^17,18,20^. In this study, sera from hamsters that were immunized at 10 μg and 0.2 μg doses as 1- and 2-dose regimens demonstrated dose-dependent binding, neutralizing and hACE2 inhibiting antibody activity, with the highest responses seen in the 2-dose regimens. The strongest responses against WA1, B.1.1.7 and B.1.351 were observed in the 10 μg group using a 2-dose regimen, with similar binding responses observed between WA1 and B.1.1.7, that were reduced slightly against B.1.351 in the 10 μg, 2-dose regimen but more substantially in the 1-dose regimens. Humoral immunity waned from peak responses at study weeks 6 and 8 to the time of challenge at week 11. Opsonizing IgG responses, shown to be associated with protection against SARS-CoV-2 in small-animal models, were also elicited^30–34^. Additionally, prior studies have demonstrated that vaccine-induced antibody Fc-mediated functions are also associated with protection against SARS-CoV-2^35–37^. Here, we demonstrated that the SpFN vaccine was able to elicit high IgG opsonization against both wild type and VOCs in hamsters, suggesting that SpFN-ALFQ vaccine induced antibodies that could leverage Fc-mediated functions associated with protection from SARS-CoV-2 VOCs to include B.1.351. Of note, we observed strong neutralizing responses in the 10 μg, 2-dose regimen groups that were similar across WA1, B.1.1.7 and B.1.351 variants. This is particularly encouraging as many studies have demonstrated decreased neutralizing antibody responses of convalescent or vaccinee sera, in particular against B.1.35^4,11,14,16,27^. hACE2-RBD binding inhibition levels mirrored overall humoral responses. This assay has functional implications for cellular entry and in conjunction with the binding and neutralization assays, will be critical to utilize in future studies to understand the critical correlates of protection conferred during SpFN-ALFQ vaccination in SGH and NHP models.

As expected from the humoral responses, mitigation of body weight loss was also observed in vaccinated animals, following SARS-CoV-2 VOC challenge, most dramatically in the 2-dose vaccinated animals. Hamsters were challenged with doses of B.1.1.7 or B.1.351 targeted to result in a 10-15% loss of body weight in control animals. In the 2-dose regimen at both 10 μg and 0.2 μg doses for B.1.1.7, loss of bodyweight was dramatically reduced, from ~12% observed in the PBS control animals to between 2-3% in the vaccinated groups. Notably, a single vaccination in the 10 μg dose group was also sufficient to confer similar protection, with the even low-dose conferring intermediate levels of protection after a single immunization. In the case of B.1.351, protection in the prime-boost regimen had a similar range of protection, from the ~13% bodyweight loss observed in the PBS treated animals to 3-4%. Single-doses were less protective than observed for B.1.1.7 challenge, but still conferred intermediate protection against B.1.351 challenge. These results are highly encouraging, showing potential protective efficacy of the SpFN-ALFQ vaccine against clinical disease caused by VOC to include B.1.351, particularly using a 2-dose regimen.

Following the challenge phase of the study, assessment of viral loads by TCID50 in lung tissues and nasal turbinate demonstrated clear protection from challenge with either B.1.1.7 or B.1.351, with complete elimination of recoverable virus detected in the majority of animals, regardless of vaccine regimen, by day 6 post challenge. Adjunctive analysis of oral swab viral load, either total RNA or subgenomic mRNA, products of discontinuous SARS-CoV-2 transcription only seen during active viral replication, confirmed the lung and nasal turbinate tissue viral culture data in showing virologic evidence of SpFN-ALFQ protection with either B.1.1.7 or B.1.351 challenge.

A key question that this study was designed to address through the utilization of SGH as a pathogenic model of SARS-CoV-2 disease, was whether SpFN-ALFQ had an impact on lung pathology developed during acute infection with B.1.1.7 or B.1.351. Challenge doses of B.1.1.7 and B.1.351 utilized in this study gave nearly equivalent pathology in unvaccinated (PBS) control animals, with significant type II pneumocyte hyperplasia and cellular infiltrate occurring within the lung on H&E, and viral staining observed by detection of viral nucleocapsid (N) protein by immunohistochemistry. For B.1.1.7 challenged animals, pathology was dramatically reduced in both the high and low-dose prime-boost groups. 1-dose 10 μg and 0.2 μg vaccine regimens conferred intermediate levels of protection, with wider variability in individual pathology within individuals. In the B.1.351 challenged animals, the 10 μg 2-dose vaccine regimen also demonstrated decreases in pathology compared with the cognate PBS control, with intermediate levels of protection conferred to the 0.2 μg groups. Taken together, SpFN-ALFQ generated robust binding antibody and neutralizing antibody responses in SGH against WA1, B.1.1.7 and B.1.351 SARS-CoV-2 variants. It was also protective in intranasal challenge with the B.1.1.7 and B.1.351 variants as assessed by body weight preservation, lung tissue viral load, oral fluid total and sgmRNA, and standard histopathological and immunohistochemical analysis of lung tissues at necropsy.

Publically available preprint reports have recently shown that the vaccines being marketed by AstraZeneca, ChAdOx1 nCoV-19 (AZD1222), and Moderna, mRNA-1273, show protection against B.1.351 challenge in hamster and rhesus models, respectively^27,38^. Another preprint describes protection from B.1.351 challenge in human ACE2 transgenic mice vaccinated with the CureVac mRNA vaccine, CVnCoV^39^. Continued molecular epidemiologic analysis of ongoing SARS-CoV-2 infections with concomitant assessment of the impact of emerging VOCs on current and next-generation SARS-CoV-2 vaccines will be essential for achieving and maintaining pandemic control, and underpins the scientific approach to both pan-SARS-CoV-2 and pan-coronavirus vaccine research and development. To this end, SpFN-ALFQ is now being evaluated in a first in human phase I clinical trial as a platform approach toward broader application to the betacoronavirus genus and the entire family of coronaviruses.

## Methods

### Transfection, Expression and Purification of SpFN 1B-06-PL

Expi293 cells (Gibco, Cat No. A14527) were maintained and passaged as per manufacturer’s guidelines in Expi293 Expression Media (Gibco, Cat No. A1435101) at 37°C, 8% CO_2_, 120 RPM, ≥ 60% RH. Briefly, the transfection reaction consisted of 1mg of purified, cGMP sourced plasmid DNA (Aldevron, pCoV 1B-06-PL) plus 3mL of Turbo 293 Transfection Reagent (Speed Biosystems, Cat No. PXX1001) mixed in 1X PBS per liter of transfected cells. The reaction was incubated at 20-25°C for 15 minutes and 125mL/flask was rapidly and aseptically transferred to 16 single use 3L Erlenmeyer shake vented flasks (Corning, Cat no. 431252) each containing 1.125L of Expi293 cells passaged at a density of 2.0 × 10^6 cells/ml on the day of transfection. After the addition of the transfection reaction, the transfected cells were incubated at 34°C, 8% CO_2_, 120 RPM, ≥ 60% RH for 5 days. The expressed product was collected, clarified by double centrifugation at 4000 RPM, 10°C, for 30 minutes, followed by depth filtration (0.65uM + 0.45uM, Sartorius, Sartobran P, Cat No. 5235306D0-SO-V) and stored at 2-8°C for further downstream processing.

SpFN 1B-06 PL was purified, concentrated and dialyzed against 50 mM Tris, 50mM NaCl, pH 8.0 solution. Briefly, 16 L of clarified expression product was initially concentrated 6 fold using a tangential flow filtration module (500 kD MW Cutoff, mPES MiniKros, Cat No.N04-500-05). The concentrate was treated with Benzonase (EMD Millipore, Cat No.EM1.01695.0001) for 120min at 22°C before conducting a buffer exchange into 50 mM Tris, 50mM NaCl, pH 7.915. The material was loaded onto a 4.4 × 14 cm Fractogel DEAE (M) Column (EMD Millipore, Cat No.1168835000) with a bed volume of 212 mL. SpFN 1B-06 PL bound to the column and was eluted with 50 mM Tris Base, 200 mM NaCl, pH 8.018, 21.55 mS/cm. Following this, a 44 mL Capto Core 400 column (Cytiva, Cat No.17372403) was used as a polishing step to remove any potential lower MW contaminants. A final dialysis step was performed to place the product into the final formulation buffer using a 300 kD MWCO UF cartridge (Repligen mPES MiniKros, Cat No. S02-E300-05). The final purified SpFN 1B-06 PL recovery yielded 55.47 mg in a volume of 215 mL, approximately a 3.47 mg/L yield from the cell expansion and growth.

### Viral stock propagation and preparation

B.1.1.7 viral stocks were generated from seed stock (USA/CA_CDC_5574/2020), obtained from BEI resources (Cat # NR-54011, Lot # 70041598) and expanded in Calu-3 cells (incubated at 37°C for 3 days). The viral stock lot used for this study (Lot # 012921-1230) was titrated in Vero-TMPRSS2 cells, with viral titers of 1.375 × 10^6^ PFU/mL. This stock was used undiluted for viral challenge with 100μL intranasally, for a final challenge concentration of 1.375 × 10^5^ PFU/mL per dose.

B.1.351 viral stocks were propagated and characterized by deep sequencing as previously described^38^. Briefly, hCoV-19/USA/MD-HP01542/2021 (B.1.1.351) was derived from the seed stock (Lot# MD-HP JHU P2) and propagated in in VeroE6-TMPRSS2 cells. The viral titer of the B.1.351 stock used for this study was 3 × 10^7^ PFU/ml in VeroE6-TMPRSS2 cells, diluted 1:100 in PBS for a challenge dose of 3×10^4^ PFU in 100μL.

### Syrian golden hamster immunizations

Male and female Syrian golden hamsters (6-8 week-old, n = 55) were acquired from Charles River Laboratories and housed at Bioqual, Inc., for the duration of the study. Following one week of acclimatization, animals were immunized intramuscularly in caudal thighs with PBS (control) or SpFN immunogen of differing doses (10 μg or 0.2 μg) pre-formulated with a fixed dose of ALFQ (20 μg of 3D-PHAD (monophosphoryl 3-deacyl lipid A (synthetic)) and 10 μg of QS21)^40^. The design, production, stability, and initial characterization of SpFN adjuvanted with ALFQ has been described previously ^20^. Briefly, the immunogen is based on the Spike protein of the WA1 SARS-CoV-2 variant with S-2P amino acid modifications expressed as a fusion protein with H. pylori ferritin in mammalian cells that self-assembles into an ordered nanoparticle each of which presents 8 Spike trimers. SpFN was formulated with ALFQ prior to administration. SpFN was produced at the WRAIR Pilot Bioproduction Facility in October 2020 as an engineering batch and stored at 4°C for 3-4 months for the prime and boost/single immunization doses, respectively.

Boosting immunizations were injected contralaterally from the prime four weeks following the prime. In-life blood sampling was conducted by retro-orbital bleeds. Maximum blood collections were determined based on animal weight and frequency of collection, in consultation with veterinary guidance. Following collections, animals were monitored until fully recovered from the anesthetic and the procedure.

### SARS-CoV-2 Syrian golden hamster challenge

Animals were challenged with SARS-CoV-2 seven weeks following the boost or single immunization. Animals were anesthetized with ketamine/xylazine and challenged by intranasal inoculation of 50 μL virus in each nostril in a drop-wise manner (100μL/hamster). Challenge dose was pre-determined for each VOC viral stock to achieve comparable clinical disease as manifested by body weight loss of 10-15%. B.1.1.7 virus was administered at a challenge dose of 1.375 × 10^5^. B.1.351 virus was administered at a challenge dose of 3 × 10^4^ PFU per hamster. Following viral challenge, all animals were weighted and observed twice daily for clinical signs (ruffled fur, hunched posture, behavior, etc.), with euthanasia criteria of 20% loss of pre-challenge body weight or becoming moribund. At study termination 6 DPC, all animals were terminally anesthetized by ketamine/xylazine, followed by exsanguination by cardiac puncture (for terminal blood collection) and euthanasia. Tissues were collected for use in downstream virologic, immunologic, molecular and histopathology studies. Animals were housed in BLS-2 during the vaccination phase and BSL-3 facilities during the challenge phase.

### Enzyme Linked Immunosorbent Assay (ELISA)

96-well Immulon “U” Bottom plates were coated with 1 μg/mL of RBD or S protein (S-2P) antigen in Dulbecco’s PBS, pH 7.4. Plates were incubated at 4°C overnight and blocked with blocking buffer (PBS containing 0.5% Casein and 0.5% BSA, pH 7.4), at room temperature (RT) for 2 h. Individual serum samples were serially diluted 2-fold in blocking buffer and added to triplicate wells and the plates were incubated for 1 hour at RT. The plates were washed with PBS containing 0.1% Tween 20, pH 7.4, followed by the addition of horseradish peroxidase (HRP)-conjugated goat anti-Hamster IgG (H+L) (1:2000 dilution) (SouthernBiotech) for an hour at RT. The HRP substrate, 2,2’-Azinobis [3-ethylbenzothiazoline-6-sulfonic acid]-diammonium salt (ABTS) (KPL) was added to the plates for 1 hour at RT. The reaction was stopped by the addition of 1% SDS per well and the absorbance (A) was measured at 405 nm (A405) using an ELISA reader Spectramax (Molecular Devices, San Jose, CA) within 30 min of stopping the reaction. The results are expressed as end point titers, defined as the reciprocal dilution that gives an absorbance value that equals twice the background value (antigen-coated wells that did not contain the test sera, but had all other components added).

### Biolayer Interferometry RBD binding assay

All biosensors were hydrated in PBS prior to use. All assay steps were performed at 30°C with agitation set at 1,000 rpm in the Octet RED96 instrument (FortéBio). HIS1K biosensors (FortéBio) were equilibrated in assay buffer (PBS) for 30 seconds before loading of His-tagged SARS-CoV-2 WA1 RBD or VOC RBDs B1.1.7, and B.1.351 (30 μg/mL diluted in PBS) for 120 seconds. Immobilized RBD proteins were then dipped in hamster sera (100x dilution with PBS) for 180 seconds followed by dissociation for 60 seconds. Binding response values were recorded at 180 seconds. SARS-CoV-2 RBD constructs (residues 331 - 527), modified to incorporate a N-terminal hexa-histadine tag (for purification), were derived from the Wuhan-Hu-1 strain genome sequence (GenBank MN9089473) and synthesized and subcloned into a CMVR plasmid by Genscript. RBD with VOC point mutations were generated using a modified QuikChange site-directed mutagenesis protocol (Agilent). The constructs resulting from site-directed mutagenesis were amplified and isolated from E. coli Stbl3 or Top10 cells. Large-scale DNA isolation was performed using either endo free Maxiprep, Megaprep or Gigaprep kits (Qiagen). All expression plasmids were transiently transfected into Expi293F cells (Thermo Fisher Scientific) using ExpiFectamine 293 transfection reagent (Thermo Fisher Scientific). Cells were grown in polycarbonate baffled shaker flasks at 34°C and 8% CO_2_ at 120 rpm. Cells were harvested 5-6 days post-transfection via centrifugation at 3,500 x g for 30 minutes. Culture supernatants were filtered with a 0.22-μm filter and stored at 4°C prior to purification. His-tagged RBD proteins were purified using Ni-NTA affinity chromatography, with 1 mL Ni-NTA resin (Thermo Scientific) used to purify protein from 1L of expression supernatant. Ni-NTA resin was equilibrated with 5 column volumes (CV) of phosphate buffered saline (PBS) (pH 7.4) followed by supernatant loading 2 x at 4°C. Unbound protein was removed by washing with 200 CV of PBS, followed by 50 CV 10mM imidazole in PBS. Bound protein was eluted with 220 mM imidazole in PBS. All proteins were further purified by size-exclusion chromatography using a 16/60 Superdex-200 purification column. Purification purity for all the proteins was assessed by SDS-PAGE.

### Biolayer Interferometry hACE2 competition assay

All biosensors were hydrated in PBS prior to use. All assay steps were performed at 30°C with agitation set at 1,000 rpm in the Octet RED96 instrument (FortéBio). SARS-CoV-2 RBD - hACE2 competition assays were carried out as follows. SARS-CoV-2 RBD (WA1 strain, 30 μg/ml diluted in PBS) was immobilized on HIS1K biosensors (FortéBio) for 180 seconds followed by baseline equilibration for 30 seconds. Serum binding was allowed to occur for 180 seconds followed by baseline equilibration (30 seconds). Recombinant hACE2 protein (30 μg/ml) was then allowed to bind for 120 seconds. Percent inhibition (PI) of hACE2 binding to the RBD by serum was determined using the equation: PI = 100 – ((hACE2 binding in the presence of mouse serum/ hACE2 binding in the absence of mouse serum) × 100).

### IgG Opsonization Assays

SARS-CoV-2 S-expressing expi293F cells were generated by transfection with linearized plasmid (pcDNA3.1) encoding codon-optimized full-length SARS-CoV-2 S protein matching the amino acid sequence of the IL-CDC-IL1/2020 isolate (GenBank ACC# MN988713), the B.1.1.7 isolate^9^, or the B.1.351 isolate ^41^. Stable transfectants were single-cell sorted and selected to obtain a high-level Spike surface expressing clone (293F-Spike-S2A). 293F-Spike-S2A cells were incubated with 100 μl of plasma diluted 100-fold in RPMI containing 10% FBS (R10) for 30 minutes at 37°C. Cells were washed 3 times and stained with a goat anti-hamster IgG (H+L) Alexa Fluor 488 (ThermoFisher Scientific). Cells were then fixed with 4% formaldehyde solution and fluorescence was evaluated on a LSRII analytic cytometer (BD Bioscience).

### SARS-CoV-1 and SARS-CoV-2 pseudovirus neutralization assay

Pseudovirions were produced by co-transfection of HEK293T/17 cells with either the SARS-CoV-1 (Sino 1-11, GenBank # AY485277) or SARS-CoV-2 (WA1/2020 GenBank # MT246667) S expression plasmid and an HIV-1 pNL4-3 luciferase reporter plasmid (pNL4-3.Luc.R-E-, NIH AIDS Reagent Program). The S expression plasmid sequences for SARS-CoV-2 and SARS-CoV-1 were codon optimized and modified to remove an 18 amino acid endoplasmic reticulum retention signal in the cytoplasmic tail in the case of SARS-CoV-2, and a 28 amino acid deletion in the cytoplasmic tail in the case of SARS-CoV-1 to improve S incorporation into pseudovirions and improve infectivity. S expression plasmids for SARS-CoV-2 VOC were similarly codon optimized, modified and included the following mutations: B.1.1.7 (69-70del, Y144del, N501Y, A570D, D614G, P681H, T718I, S982A, D1118H), B.1.351 (L18F, D80A, D215G, 241-243del, K417N, E484K, N501Y, D614G, A701V, E1195Q). Virions pseudotyped with the vesicular stomatitis virus (VSV) G protein were used as a non-specific control. Infectivity and neutralization titers were determined using ACE2-expressing HEK293 target cells (Integral Molecular). Test sera were diluted 1:40 in cell culture medium and serially diluted; then 25 μL/well was added to a white 96-well plate. An equal volume of diluted SARS-CoV-2 PSV was added to each well and plates were incubated for 1 hour at 37°C. Target cells were added to each well (40,000 cells/ well) and plates were incubated for an additional 48 hours. Relative light units (RLU) were measured with the EnVision Multimode Plate Reader (Perkin Elmer, Waltham, MA) using the Bright-Glo Luciferase Assay System (Promega, Madison, WI). Neutralization dose–response curves were fitted by nonlinear regression using the LabKey Server. Final titers are reported as the reciprocal of the dilution of serum necessary to achieve 50% (ID_50_, 50% inhibitory dose). Assay equivalency for SARS-CoV-2 was established by participation in the SARS-CoV-2 Neutralizing Assay Concordance Survey (SNACS) run by the Virology Quality Assurance Program and External Quality Assurance Program Oversite Laboratory (EQAPOL) at the Duke Human Vaccine Institute, sponsored through programs supported by the National Institute of Allergy and Infectious Diseases, Division of AIDS.

### Oral cavity viral RNA measurements

The oral cavity was swabbed with a sterile flocked swab that was immediately placed into a cryovial with 1mL PBS. Vials were snap-frozen on dry ice and stored at −80°C until testing. Viral RNA was extracted from 200 ul of oral swab material using the Qiagen EZ1 DSP Virus kit on the automated EZ1 XL Advance instrument (Qiagen, Valencia, CA). Real-time quantitative reverse transcription – polymerase chain reactions (RT-qPCR) were performed on the 7500 Dx Fast thermal cycler (Thermo Fisher Scientific, Life Technologies, Carlsbad, CA). SARS-CoV-2 specific forward and reverse primers and probe targeting the E gene encoding the envelope protein were used for amplification of the viral RNA. Amplification of the sgmRNA was achieved using the Leader TRS sequence specific primer, the reverse E primer and the E specific probe as described previously^18^. A synthetic RNA for subgenomic E was used as a calibrator. Final results were reported in copies/ml.

### Tissue viral burden by TCID50

The infectious titer determination from lungs and nasal turbinates was obtained by performing a TCID50 assay. Vero TMPRSS2 cells were plated at 25,000 cells per well in DMEM supplemented with 10% FBS and gentamicin. Cells were incubated at 37°C, 5.0% CO_2_. When cells achieved 80-100% confluency, the media was aspirated and replaced with 180μL of DMEM containing 2% FBS and gentamicin. 0.10-0.20 mg sections of the right lobe of the lung and the nasal turbinates were collected at necropsy, snap frozen, and stored at −80°C until processing. Frozen tissue was placed in 15 mL conical tube on wet ice containing 0.5 mL media and homogenized 10-30 secs (Probe, Omni International: 32750H). The tissue homogenate was spun to remove debris at 2000 *g*, 4°C for 10 min. The supernatant was then passed through a strainer (Pluriselect: Cat No. 43-10100-40) into the original vial and kept on wet ice. From the strained supernatant, 20 μL aliquots were tested in quadruplicate in a 96-well plate format. The top row of the 96-well plate was mixed 5 times and serially diluted by pipette transfer of 20 μL, representing 10-fold dilutions. Pipette tips were disposed of between each row and mixing was repeated until the last row on the plate. After incubation for 4 days, wells were visually inspected for cytopathic effects (CPE) scored as CPE minus (-) where non-infected wells have a clear confluent cell layer, or CPE plus (+), where rounding of infected cells is observed. Positive controls were utilized for optimal assay performance, where the TCID50 tested within 2-fold of the expected value. TCID50 of all samples were calculated using the Read-Muench formula.

### Histology and immunohistochemistry

Lungs from 6 DPC were insufflated and perfused with 10% neutral-buffered formalin. Three tissue sections from each of the left lung lobes were used to evaluate the lung pathology. Sections were processed routinely into paraffin wax, then sectioned at 5 μm, and resulting slides were stained with hematoxylin and eosin. Immunohistochemistry (IHC) on formalin fixed paraffin embedded tissue sections was performed using the Dako Envision system (Dako Agilent Pathology Solutions, Carpinteria, CA, USA). Briefly, after deparaffinization, peroxidase blocking, and antigen retrieval, sections were stained with an anti-SARS-CoV/SARS-COV-2 nucleocapsid (N) protein rabbit monoclonal antibody (#40143-R001, Sino Biological, Chesterbrook, PA, USA) at a dilution of 1:6000 and incubated at RT for 45 min. Sections were rinsed and stained with peroxidase-labeled polymer (secondary antibody) for 30 min. Slides were rinsed and a brown chromogenic substrate 3,3’ Diaminobenzidine (DAB) solution (Dako Agilent Pathology Solutions) was applied for 8 min. Slides were rinsed, counterstained with hematoxylin, and rinsed. The sections were dehydrated, cleared with Xyless II, and cover-slipped. All tissue slides were evaluated by a board-certified veterinary anatomic pathologist blinded to study group allocations. Semi-quantitative scoring of pulmonary pathology was performed, with grading of intra-alveolar edema, type II pneumocyte hyperplasia, mononuclear cellular infiltrates, polymorphonuclear cellular infiltrates, alveolar histiocytosis, alveolar necrosis, bronchioalveolar epithelial degeneration, bronchiolar epithelial hyperplasia, and interstitial collagenous deposition. Each finding was scored as follows: 0 - absent, 1 - minimal (<10% of tissue section affected); 2 - mild (11-25% of tissue section affected); 3 - moderate (26-50% of tissue section affected); 4 - marked (51-75% affected); 5- severe (>75% of tissue section affected). IHC sections were examined at 400X magnification and evaluated for the number of immunopositive cells per slide.

### Ethical Statement

All animal *in vivo* procedures were carried out in accordance with institutional, local, state, and national guidelines and laws governing research in animals including the Animal Welfare Act. Animal protocols and procedures were reviewed and approved by the Animal Care and Use Committee of both the US Army Medical Research and Development Command (USAMRDC) Animal Care and Use Review Office as well as the Institutional Animal Care and Use Committee of Bioqual, Inc. (protocol number 20-144). Bioqual, Inc. and the USAMRDC are accredited by the Association for Assessment and Accreditation of Laboratory Animal Care and are in full compliance with the Animal Welfare Act and Public Health Service Policy on Humane Care and Use of Laboratory Animals. Oversight of all research was approved and conducted by the WRAIR Institutional Biological Safety Committee.

### Statistical Analysis

All statistical analysis were performed using GraphPad Prism version 8 software. All statistical tests shown were performed using the Kruskal-Wallis test with Dunn’s correction, comparing all variables (multiple comparison analysis, not against a standard control). Actual p-values for each comparison annotated for statistically relevant values, or for values near statistical significance. All other not statistically or biologically relevant values reported as not-significant (ns). Comparisons to PBS control group are shown directly above each vaccine treatment condition (gray), with inter/intra vaccine regimen comparisons indicated by a solid line between groups, with the statistical result directly above (black).

## Acknowledgments

We thank Morgane Rolland for molecular phylogeny and Misook Choe for SpFN expression consultation. We also thank Sebastian Molnar, Erin Kavusak, Jonah Heller, Claudelle Busano, Hannah King, Sylriel Peters, Theron Jenifer and Jean-Paul Todd for technical support. We also thank Mihret Amare, Mekdi Taddese, Jarrett Headley and Yahel Romem for programmatic support and planning and Paul Scott scientific inputs and administrative oversight. This work was funded by the US Department of Defense, Defense Health Agency was executed, in part, through a cooperative agreement (W81XWH-18-2-0040) between the Henry M. Jackson Foundation for the Advancement of Military Medicine, Inc., and the U.S. Department of Defense (DOD). Material has been reviewed by the Walter Reed Army Institute of Research. The opinions or assertions contained herein are the private views of the authors, and are not to be construed as official, or as reflecting true views of the Department of the Army or the Department of Defense.

## Author contributions

Conceptualization, D.L.B., M.G.J., N.L.M., K.M.; Investigation, K.MW., E.K.B, W-H.C., E.J.M., I.L-N., L. L.J., D.P-P., G.D.G, I.S., A.G., C.K., S.S., H.He., H.G., H.Ha., S.Ka., M.P., A.W., K.R., X.Z, M.R., S.A.P, G.D.G, E.K.B., D.L.B., M.G.J., N.L.M., K.M.; Data Curation—K.MW., D.L.B., M.G.J. Essential Reagents, R.S.S., A.A., E.B.M., P.M., S.J.K., A.S.P., J.R.C., M.S., G.R.M., E.A.B., D.C.D., R.A.S., Writing – Original Draft, K.MW, D.L.B., N.L.M., K.M.; Writing – Review & Editing, All authors; Visualization—K.MW., E.K.B., E.J.M., P.V.T, D.L.B., M.G.J., N.L.M. K.M.; Supervision—D.L.B., M. G.J., S.A.P., J.D., S.P.D., J.W.F., M.G.L., S.V., N.L.M., K.M.; Funding Acquisition—K.M., N.L.M.

## Supplemental material

**Supplemental Figure 1. Additional antibody responses following SpFN-ALFQ immunization**. **a**, ELISA was performed using either WA1 derived Receptor Binding Domain (RBD) or S-2P Spike proteins from sera taken at week 8. The vaccine regimens are indicated on the x-axis by phosphate buffered saline (PBS) control or SpFN dose with the number of vaccinations in the regimen given parenthetically and by color code (blue, 2-dose, red, 1-dose). Endpoint titers are given on the y-axis as geometric mean titers with data displayed in box plots with the top and bottom bars of the box the standard deviation and the middle bar as the median value. P-values for SpFN-ALFQ vaccination groups compared with PBS control are given just above the boxes in light grey while inter- and intra-regimen p-values are given above the boxes in black. ns, not significant (p > 0.05). **b**, Octet Biolayer Interferometry (BLI) responses against the WA1, B.1.1.7, and B.1.351 sequences of the RBD are given for the vaccination regimens as in **a** on the x-axis from sera collected at week 8. BLI responses are given in nanometers (nm) on the y-axis. Vaccination regimen color coding and statistical treatments are as in **a**. **c**, IgG opsonization as measured by binding to SARS-CoV-2 Spike protein expressing expi293F cells subsequently stained by fluorescently tagged goat anti-hamster IgG and detected by flow cytometry. Fluorescence is given as mean fluorescence intensity (MFI) on the y-axis. Vaccination regiment color coding and statistical treatments are as in **a**.

**Supplemental Fig 2. Quantitative SARS-CoV-2 total and subgenomic mRNA (sgmRNA) viral load from oral swabs following challenge**. Oral fluid collection from were obtained by swabbing at days 2, 4, and 6 post challenge in hamsters and submitted to quantitative SARS-CoV-2 total and sgmRNA analysis. Viral load is given on y-axes at copies/mL of oral fluid plotted on a log10 scale. Vaccination regimen groups, stratified by day post-challenge, are given on the x-axes: Phosphate buffered saline (PBS) control, gray circles; 10 μg and 0.2 μg 2-dose regimen groups, blue circles; 10 μg and 0.2 μg 1-dose regimen groups, red circles. Data are given in box plots where the horizontal bar is the median value for each group. Where values between groups are statistically meaningfully different, p-values are given above horizontal black bars showing the analyzed groups. In all other cases, the p-values were > 0.05. **a**, sgmRNA viral load from B.1.1.7 challenged hamsters; (B) total RNA viral load from B.1.1.7 challenged hamsters; **c**, sgmRNA viral load from B.1.351 challenged hamsters; **d**, total RNA viral load from B.1. 351 challenged hamsters.

**Supplemental Table 1. Semi-quantitative histopathology and IHC scores**

Lung tissues were collected at necropsy on day 6 post-challenge, fixed with neutral buffered formalin, and stained with hematoxylin and eosin (H&E) for standard microscopic examination as well as submitted for immunohistochemical (IHC) staining for SARS-CoV-2 nucleocapsid (N) protein. (A) H&E sections were semi-quantitatively scored on the histopathology present in all lung sections examined (Methods). The overall severity of interstitial pneumonia (IP) present in individual animals in the PBS control and vaccinated groups is given. (B) IHC sections were examined at 400X magnification, and evaluated for the number of immunopositive cells per slide. Distribution of the number of immunopositive cells per section is given per PBS control and 2-dose and 1-dose vaccine group. The number of vaccinations per vaccine group are given parenthetically.

## Notes

### Competing Interest Statement

Drs. Modjarrad and Joyce are co-inventors on a pending patent application for the Spike Protein Ferritin Nanoparticle (PCT/US2021/021405) Assigned to the U.S. Government as represented by the Secretary of the Army.

