## Supplemental Materials for "A SARS-CoV-2 spike ferritin nanoparticle vaccine protects against heterologous challenge with B.1.1.7 and B.1.351 virus variants in Syrian golden hamsters"

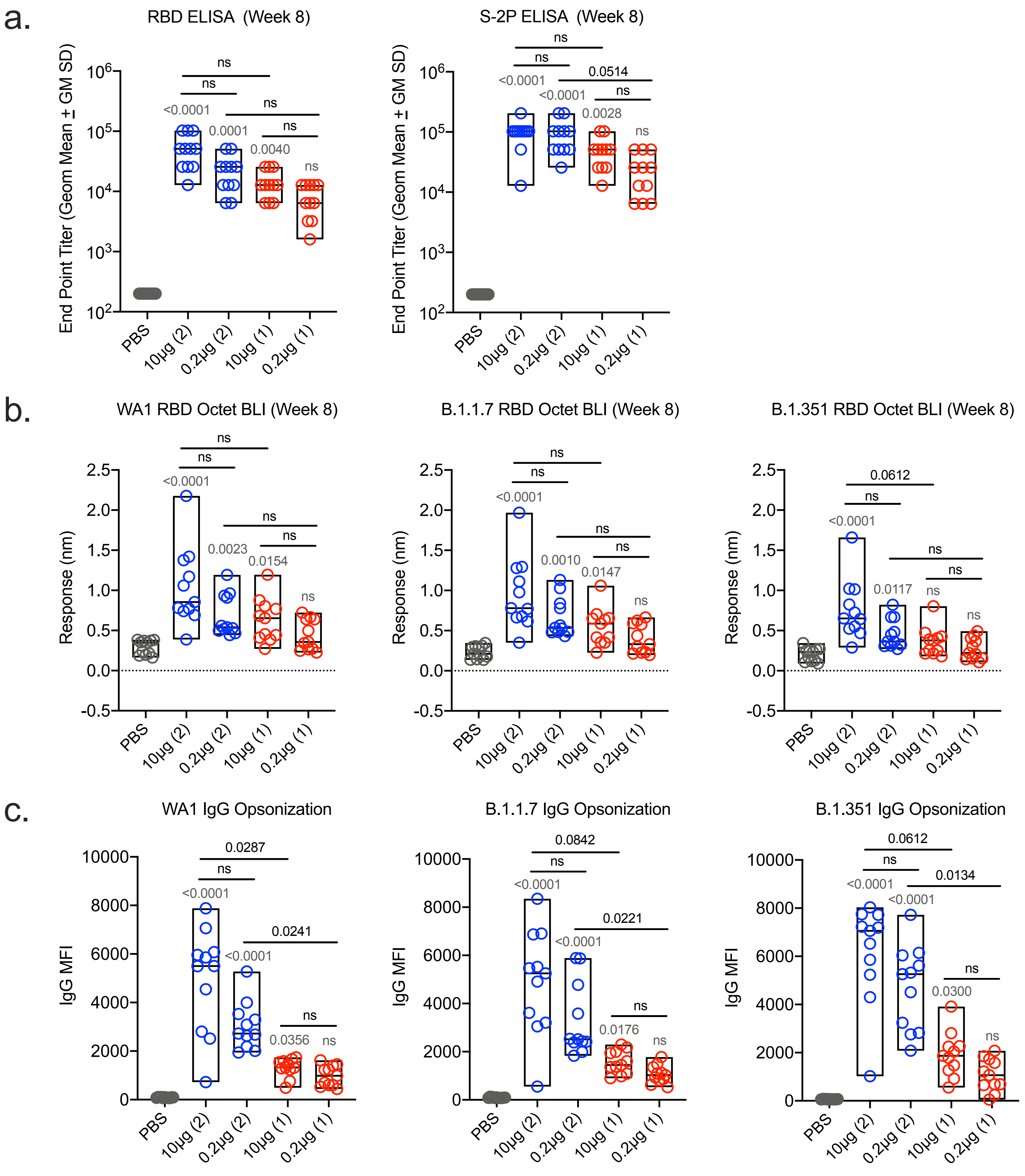
a.
RBD ELISA (Week 8)
S-2P ELISA (Week 8)
End Point Titer (Geom Mean ± GM SD)
<0.0001
0.0001
0.0040
<0.0001
0.0028
0.0514
ns
PBS
10μg (2)
0.2μg (2)
10μg (1)
0.2μg (1)
b.
WA1 RBD Octet BLI (Week 8)
B.1.1.7 RBD Octet BLI (Week 8)
B.1.351 RBD Octet BLI (Week 8)
Response (nm)
0.0023
0.0154
0.0010
0.0147
0.0117
0.0612
c.
WA1 IgG Opsonization
B.1.1.7 IgG Opsonization
B.1.351 IgG Opsonization
IgG MFI
0.0287
0.0241
0.0356
0.0842
0.0221
0.0176
0.0134
0.0300

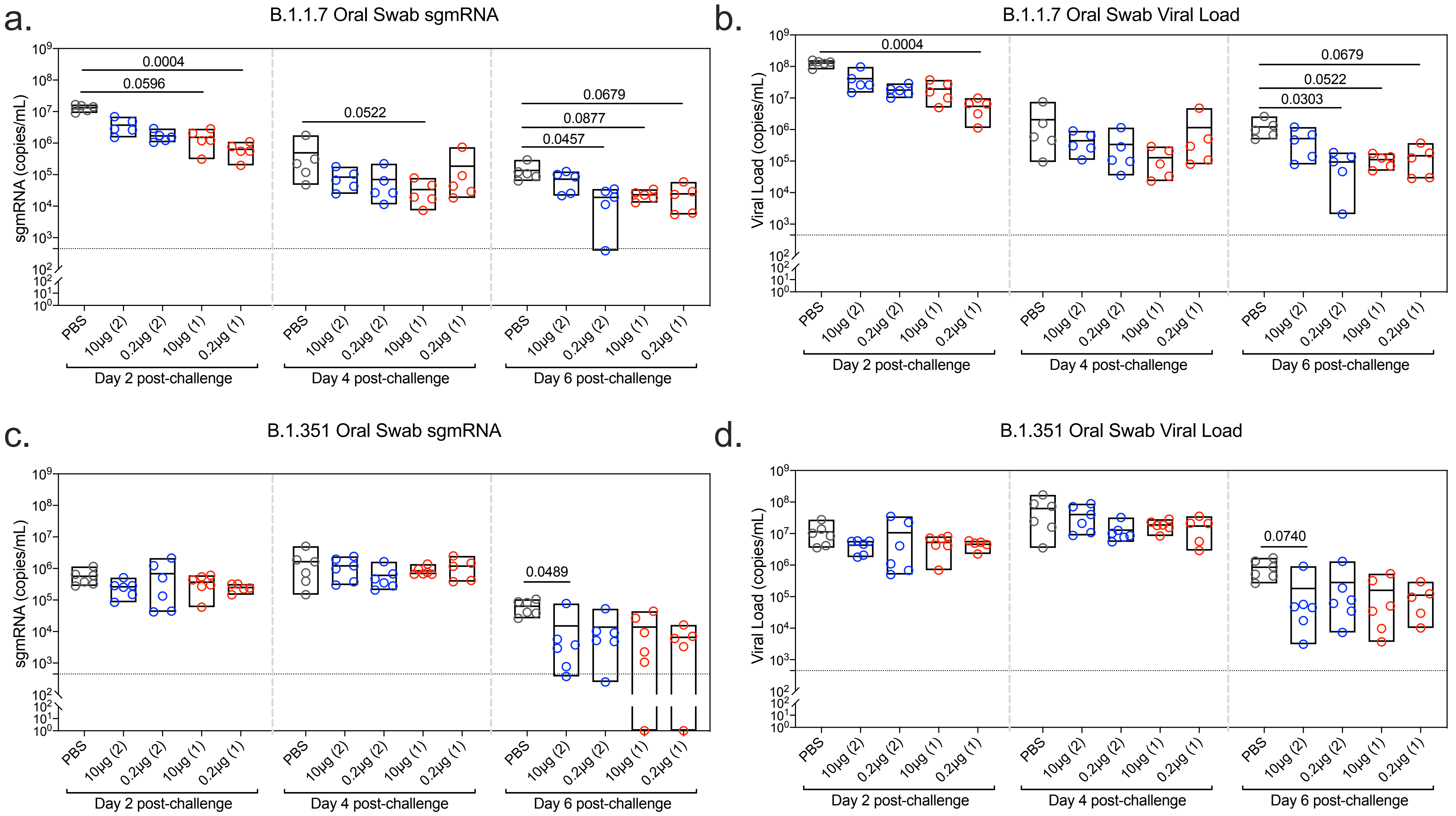

a.
B.1.1.7 Oral Swab sgmRNA
sgmRNA (copies/mL)
0.0004
0.0596
0.0522
0.0679
0.0877
0.0457
PBS
10μg (2)
0.2μg (2)
10μg (1)
0.2μg (1)
Day 2 post-challenge
Day 4 post-challenge
Day 6 post-challenge
b.
B.1.1.7 Oral Swab Viral Load
Viral Load (copies/mL)
0.0004
0.0679
0.0522
0.0303
c.
B.1.351 Oral Swab sgmRNA
sgmRNA (copies/mL)
0.0489
d.
B.1.351 Oral Swab Viral Load
Viral Load (copies/mL)
0.0740

### Supplemental Table 1A

| Challenge Strain | Vaccine regimen | Vaccine dose | IP Pathology (# observed/total n) | | | | |
|---|---|---|---|---|---|---|---|
| | | | None | Minimal | Mild | Moderate | Marked |
| B.1.1.7 | Prime-boost | PBS | | | | 3/5 | 2/5 |
| | | 10 µg (2) | 3/5 | 2/5 | | | |
| | | 0.2 µg (2) | 2/5 | 3/5 | | | |
| | Single | 10 µg (1) | 2/5 | 1/5 | 1/5 | | 1/5 |
| | | 0.2 µg (1) | 1/5 | 1/5 | | 2/5 | 1/5 |
| B.1.351 | Prime-boost | PBS | | | | 4/6 | 2/6 |
| | | 10 µg (2) | 4/6 | 1/6 | 1/6 | | |
| | | 0.2 µg (2) | 1/6 | 2/6 | 1/6 | 1/6 | 1/6 |
| | Single | 10 µg (1) | 1/6 | 3/6 | 2/6 | | |
| | | 0.2 µg (1)* | | 3/5 | 1/5 | | 1/5 |

*one animal lost to study due to injury unrelated to infection.

### Supplemental Table 1B

| Challenge Strain | Vaccine regimen | Vaccine dose | SARS-CoV-2 Viral Antigen per Section (# animals observed/total n) | | | | | |
|---|---|---|---|---|---|---|---|---|
| | | | None | Single cell | <5 cells | <20 cells | <40 cells | >500 cells |
| B.1.1.7 | Prime-boost | PBS | | | | | | 5/5 |
| | | 10 µg (2) | 3/5 | 2/5 | | | | |
| | | 0.2 µg (2) | 4/5 | 1/5 | | | | |
| | Single | 10 µg (1) | 3/5 | | 1/5 | 1/5 | | |
| | | 0.2 µg (1) | 1/5 | 1/5 | 1/5 | 2/5 | | |
| B.1.351 | Prime-boost | PBS | | | | | | 6/6 |
| | | 10 µg (2) | 5/6 | | | 1/6 | | |
| | | 0.2 µg (2) | 1/6 | 5/6 | | | | |
| | Single | 10 µg (1) | 4/6 | | | 2/6 | | |
| | | 0.2 µg (1)* | | 2/5 | 2/5 | | 1/5 | |

*one animal lost to study due to injury unrelated to infection.

#### Supplemental material

**Supplemental Figure 1. Additional antibody responses following SpFN-ALFQ immunization**. **a**, ELISA was performed using either WA1 derived Receptor Binding Domain (RBD) or S-2P Spike proteins from sera taken at week 8. The vaccine regimens are indicated on the x-axis by phosphate buffered saline (PBS) control or SpFN dose with the number of vaccinations in the regimen given parenthetically and by color code (blue, 2- dose, red, 1- dose). Endpoint titers are given on the y-axis as geometric mean titers with data displayed in box plots with the top and bottom bars of the box the standard deviation and the middle bar as the median value. P-values for SpFN-ALFQ vaccination groups compared with PBS control are given just above the boxes in light grey while inter- and intra-regimen p-values are given above the boxes in black. ns, not significant ($p > 0.05$). **b**, Octet Biolayer Interferometry (BLI) responses against the WA1, B.1.1.7, and B.1.351 sequences of the RBD are given for the vaccination regimens as in **a** on the x-axis from sera collected at week 8. BLI responses are given in nanometers (nm) on the y-axis. Vaccination regimen color coding and statistical treatments are as in **a**. **c**, IgG opsonization as measured by binding to SARS-CoV-2 Spike protein expressing expi293F cells subsequently stained by fluorescently tagged goat anti-hamster IgG and detected by flow cytometry. Fluorescence is given as mean fluorescence intensity (MFI) on the y-axis. Vaccination regiment color coding and statistical treatments are as in **a**.
